# Hybridization and Low Genetic Diversity in the Endangered Alabama Red-Bellied Turtle (*Pseudemys alabamensis*)

**DOI:** 10.1101/2021.10.30.466626

**Authors:** Nickolas Moreno, Andrew Heaton, Kaylin Bruening, Emma Milligan, David Nelson, Scott Glaberman, Ylenia Chiari

**Affiliations:** Department of Biology, University of South Alabama, Mobile, AL, USA; Department of Biology, George Mason University, Fairfax, VA, USA; Grand Bay National Estuarine Research Reserve, Mississippi Department of Marine Resources, Moss Point, MS, USA; Department of Environmental Science and Policy, George Mason University, Fairfax, VA, USA

**Author notes:** **Corresponding Author:** Dr. Ylenia Chiari, George Mason University, Department of Biology, 4400 University Dr., Fairfax, VA 22030, USA.

**Keywords:** Conservation, Endemism, Microsatellites, Mitochondrial DNA, Southeastern United States, Turtles

## Abstract

*Pseudemys alabamensis* is one of the most endangered turtle species in the United States due to its small population size and restricted geographic distribution in coastal Alabama and Mississippi. Increased urbanization and climate change impacts in the region further threaten this species. Populations of *P. alabamensis* are geographically isolated from one another by land and salt water, which could act as barriers to intraspecific gene flow. It is currently unknown how differentiated these isolated populations are from one another or whether they have experienced reductions in population size. Previous work found morphological differences between Alabama and Mississippi populations, suggesting that they may be evolutionarily distinct. Other *Pseudemys* turtles such as *P. concinna* and *P. floridana* occur within the same geographic area as *P. alabamensis* and are known to hybridize with each other. These more abundant species could further threaten the unique genetic identity of *P. alabamensis* through introgression. In order to evaluate the endangered status of *P. alabamensis* and the level of hybridization with other species, we used the mitochondrial (mtDNA) control region and nuclear microsatellite markers to assess genetic variation within and among populations of this species throughout its range and estimate admixture with co-occurring *Pseudemys* species. Genetic diversity of *P. alabamensis* was lower than expected at both markers (no variation in mtDNA and excess of homozygosity in microsatellites). We found evidence of genetic differentiation between Alabama and Mississippi populations as well as two populations (Fowl River, Alabama and Biloxi, Mississippi) with low estimated breeding sizes and signs of inbreeding. Finally, we found evidence of admixture of *P. alabamensis* with *P. concinna*/*P. floridana* and *Pseudemys* peninsularis (a species not native to Alabama or Mississippi). Our results indicate that *P. alabamensis* is highly endangered throughout its range and threatened by both low population sizes and hybridization. In order to improve the species’ chances of survival, focus should be placed on habitat preservation, maintenance of genetic diversity within both Mississippi and Alabama populations, and regular population monitoring activities such as nest surveillance and estimates of recruitment.

## INTRODUCTION

The southeastern United States is a biodiversity hot-spot, harboring higher levels of endemic species than other areas of the country (Jenkins et al. 2015). Alabama, in particular, has a high concentration of regionally endemic species, especially freshwater turtles, and is within one of three global turtle priority areas for conservation (Buhlmann et al. 2009, Lydeard and Mayden 1995). Freshwater turtles are a conservation concern worldwide, with >60% of species classified as threatened (Buhlmann et al. 2009). While some turtle species in the southeastern US are not currently imperiled, others have multiple risk factors for extinction such as low population size and restricted habitat range (IUCN 2001, Mace et al 2008, Purvis et al. 2000). The Alabama red-bellied turtle (*Pseudemys alabamensis*) is among the most at-risk species in the U.S. and considered by some authors as “the most endangered turtle on the continent” (Spinks et al. 2013). Although it is classified as endangered by both the U.S. Fish and Wildlife Service (USFW 1987) and the International Union for Conservation of Nature (IUCN) Red List, studies on this species across its entire distribution are lacking. This dearth of information prevents development of targeted management and conservation actions. Although *P. alabamensis* does occur within some protected areas (Heaton et al. in press), there are currently no specific survey activities or targeted management actions to ensure monitoring and protection of this species.

*Pseudemys alabamensis* is threatened by habitat modification, including dredging, road-kill of adults and juveniles, and also competition with other species (Nelson et al. 2009). Turtles may also be used for shooting practice (Alexander, 2018). This species also occurs within a very limited distribution range and is found exclusively in coastal rivers along Mobile Bay in Alabama and the Mississippi Sound (Fig. 1) (Leary et al., 2008). An isolated population once existed further inland in southwestern Alabama, but has since been extirpated (Mount 1975). The freshwater bodies currently inhabited by *P. alabamensis* are separated from one another by land and salt water, which likely prevents substantial movement of individuals between river populations. Some morphological differences have been previously noted between Alabama and Mississippi populations of *P. alabamensis* such as the dorsal width of the cervical scute (Leary et al. 2003), supporting the existence of isolated populations within this species.

**Figure 1.**
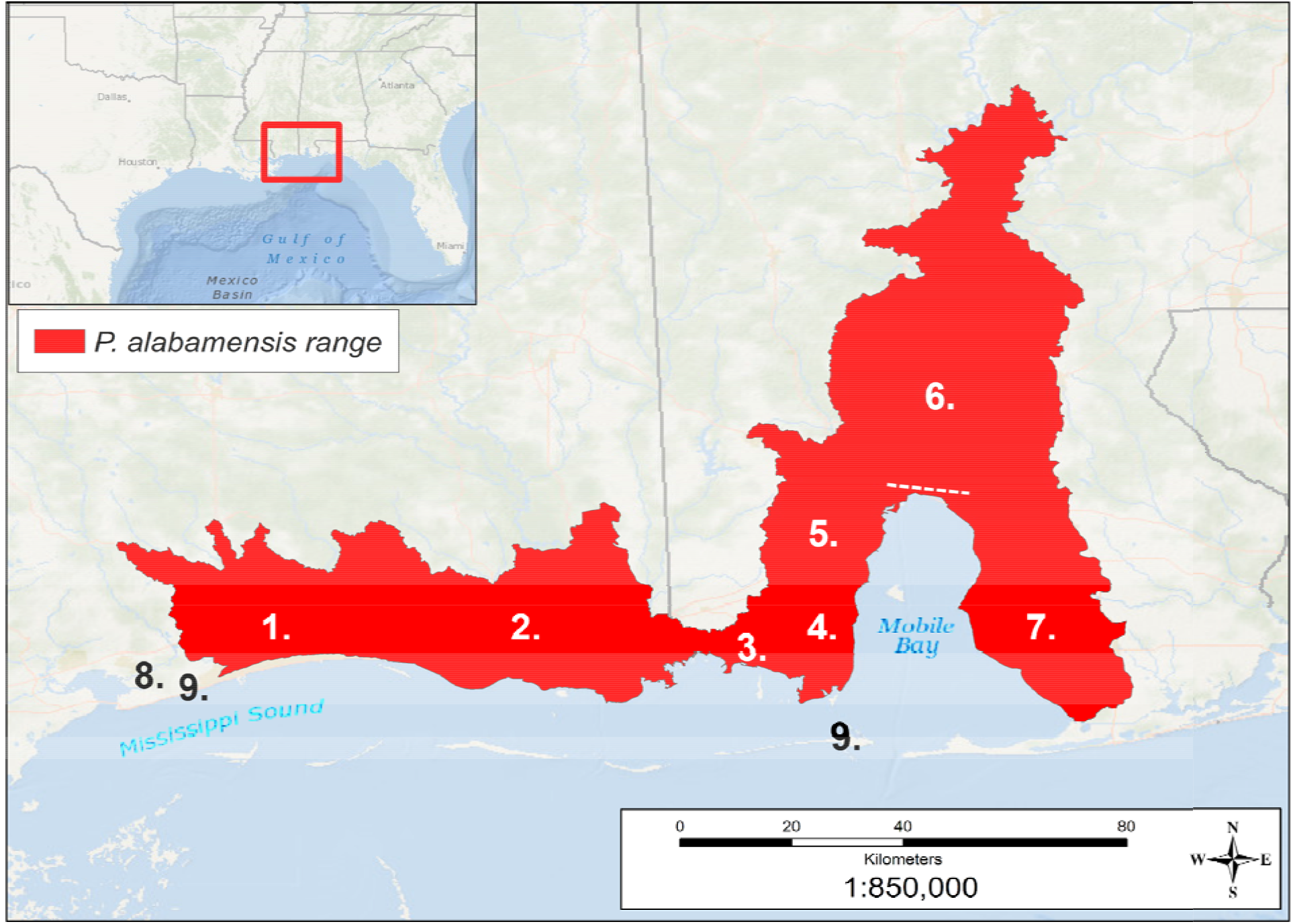
Projected Range of *P. alabamensis* based on GIS-defined hydrologic unit compartments (HUCs) created around capture locations from this study along with data from Nelson 1994, 1995, 1996, 1997, 1998, Leary et al. 2003, and Jackson et al. 2012. Approximate location of rivers sampled within the range marked by numbers as follows (numbers as in Table 1): 1. Biloxi, 2. Pascagoula, 3. Bayou La Batre, 4. Fowl River, 5. Dog River, 6. Mobile-Tensaw Delta [Mobile Bay Causeway (US HWY 98) indicated with dashed line], 7. Weeks Bay, 8. Wolf River Individual, 9. Waif Localities.

Despite the small range and fragmented populations of *P. alabamensis*, virtually nothing is known about key factors needed for developing a species survival plan such as population size, potential existence of genetically differentiated populations, and estimates of the level of admixture with closely related sympatric *Pseudemys* species. Many of these issues can be resolved with a range-wide study to assess population connectivity, genetic diversity, and levels of admixture among sympatric populations, and to establish appropriate conservation units for this species and consequently identify priority areas for monitoring and protection. To date, genetic data on *P. alabamensis* have been collected only on a relatively small sample size to clarify the taxonomic status of this species (Jackson et al. 2012, Spinks et al. 2013) or to assess genetic diversity at a single locality (Hieb et al. 2014). These studies found complex relationships between species in the *Pseudemys* genus, possibly originating from hybridization and introgression, and low genetic diversity for the Mobile-Tensaw Delta population of *P. alabamensis* in Alabama.

Although hybridization has been observed within the *Pseudemys* genus, there are no documented cases of hybridization with *P. alabamensis*, even if this species co-occurs with two other *Pseudemys* species, *P. concinna* and *P. floridana*, which are known to hybridize in the area (Mount 1975). In addition to observed hybridization of other *Pseudemys* species, we have also made anecdotal observations of mixed shell morphologies within *P. alabamensis* (Moreno pers. obs.). Introgression with native *P. concinna* and *P. floridana*, or non-native species that may have been introduced to the area, would have major conservation implications for *P. alabamensis*, as it would threaten the unique genetic identity of an already highly geographically restricted species with a likely low population size.

Here, we utilize mitochondrial DNA (mtDNA) and microsatellite markers to identify genetic structuring of populations of *P. alabamensis*, measure intraspecific genetic diversity, investigate the possibility of recent reductions in population sizes, and assess potential hybridization with sympatric species. Our results, in collaboration with local conservation organizations and authorities can directly inform monitoring and conservation activities including protecting nesting sites, assessing recruitment, identifying major threats to isolated populations, and monitoring population sizes by mark-recapture efforts. Finally, as climate change increasingly impacts coastal populations, understanding the current distribution of *P. alabamensis*, as well as predicting how changing water and salinity levels and habitat and food availability will affect this species, will be critical for determining its long-term survival potential.

## METHODS

### Permits

This research was conducted under U.S. Fish and Wildlife Service permit #TE40523A-2, Mississippi Department of Wildlife, Fisheries, and Parks permit #0614181, and Alabama Fish and Wildlife permits #2018063278468680 and #2019097050868680. Trapping and handling methods were approved by the University of South Alabama Institutional Animal Care and Use Committee (IACUC Protocol No. 921991-3).

### Sample Collection

Fieldwork was carried out from 2018-2019 throughout the range of *P. alabamensis* (Fig. 1). Trapping of *Pseudemys* turtles was performed with encounter-type aquatic hoop traps. Hoop traps were composed of an interior lead net and a double throated hoop trap attached at each end (paired net method). Hoop nets were 1.2m in diameter and 4.6m in length, while lead nets were 1.2m in height and 9-12m in length. Floats were added to hoop nets to maintain flotation and ensure access to air. Nets were anchored to the substrate with PVC tubing. Traps were left un-baited and checked once every 36 hours. Specific trap site selection was based on multiple factors: water depth, substrate, disturbance, basking logs, observed boat traffic, and submerged aquatic vegetation. In addition to trapping turtles, samples were also collected from roadkill individuals on the Mobile Bay Causeway (Fig. 1), an area known for high rates of mortality for the species. Finally, we also collected samples from two waif individuals found at Dauphin Island, Alabama and Gulfport, Mississippi. The geographic locations of sampling sites were recorded with a handheld GPS. To prevent re-sampling, turtles were marked for identification by notching the marginal scutes. Because of admixture between individuals of the cooter complex in the area (*P. concinna* and *P. floridana*), many individuals captured in this study presented mixed morphological characteristics; therefore individuals were ID’d to the most similar species following morphological descriptions of the species in Alabama as in Mount (1975) and Leary (2008). Briefly, *P. alabamensis* possesses an upper jaw with central notch flanked by a cusp on each side, complete eye bar, and and a prefrontal arrow formed from the meeting of the sagittal head stripes with the supratemporal stripes. *P. concinna* possesses a smooth upper jaw, usually possessing a marked plastron and “C”-shaped marking on plural scutes, and lacking a complete eye bar. *P. floridana* has an unmarked plastron, unmarked undersides of posterior marginal scutes, a vertical bar on pleural scutes, and complete eye bars (Fig. 2).

**Figure 2.**
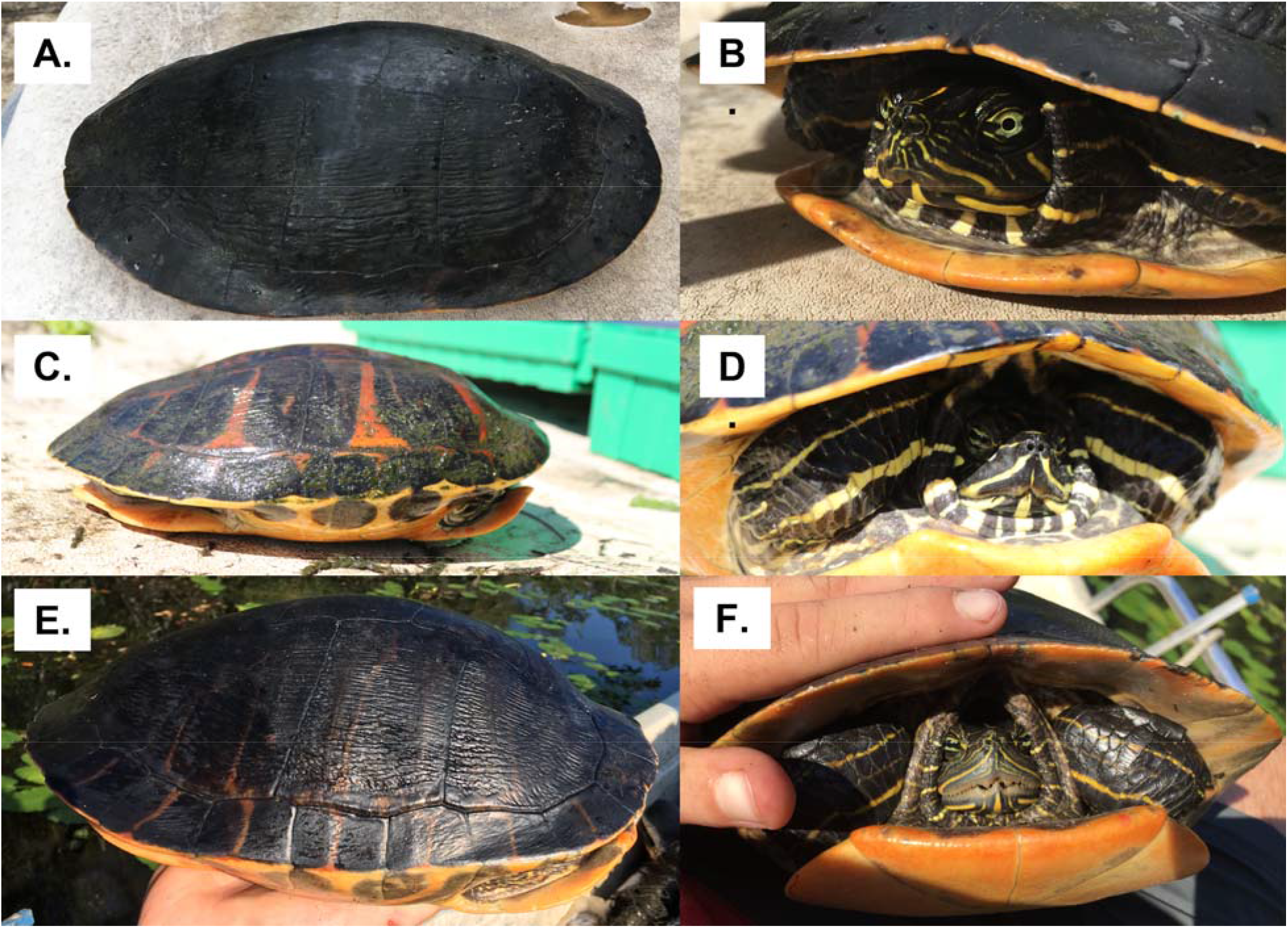
Photos illustrating two captured individual turtles that based on morphological characteristics were considered to be potential hybrids. **A & B**. Individual identified as *P. alabamensis*/*P. concinna* hybrid due to strongly reduced jaw cusp, incomplete eye bars, and incomplete prefrontal arrow. Individual was found in Bayou La Batre, Mobile County, Alabama. **C & D**. Individual considered to be *P. alabamensis*/*P. peninsularis* hybrid due to resemblance to *P. peninsularis* and presence of *P. alabamensis* characteristics. Individual found in Dog River, Mobile County, Alabama. **E & F**. *P. alabamensis* individual showing no morphological characteristics that could be considered as a sign of hybridization.

Blood for DNA extractions was collected from the subcarapacial sinus of each turtle. The skin of animals at the site was treated with 70% isopropyl alcohol prior to drawing blood. A maximum of 0.5% of body weight was collected from each animal using a 23-gauge needle and a 3-ml syringe. All animals were released at the point of capture after blood sampling was performed and after ensuring that the puncture site was not bleeding and the animal was well. Blood was stored in 2ml eppendorf tubes with 1ml of prepared blood preservative that consisted of 100mM Tris-HCL, 100mM EDTA, 10mM NaCl, and 0.5% SDS. Samples were stored on ice until returned to the lab where they were then placed at -20 C for long term storage until DNA extractions were performed. DNA extractions were carried out using the Qiagen DNeasy Blood and Tissue kit (Qiagen, Inc., Valencia, Ca) following the manufacturer’s instructions for nucleated blood.

### Mitochondrial DNA amplification and analysis

Fragments of the mitochondrial control region were amplified using the primers Des-1 and Des-2, which were originally developed by Starkey et al. (2003) for the painted turtle (Chrysemys picta). Twenty-five ul reactions were prepared using 12.5ul GoTaq G2 Green Master Mix (Promega), 0.5ul 10mg/ml bovine serum albumin, 1.2ul each of 10mM forward and reverse primers, 6.8ul H2O, and 2.8ul DNA extract. PCR conditions were as follows: 95°C for 3 min, 35 cycles of 95°C for 1 min, 55°C for 30 s, 72°C for 1 min; and a final 10 min extension at 72°C. PCR products were checked on a 1% agarose gel to ensure proper amplification and then purified using Exosap-It (Applied Biosystems) according to manufacturer’s instructions. Sequencing was carried out by the DNA Analysis Facility at Yale University. Sequences were checked and manually edited when necessary using FinchTV (Treves 2010). Cleaned sequences were aligned and collapsed into haplotypes using UGENE (Okonechnikov et al. 2012). Haplotypes were inputted into a BLAST (Basic Local Alignment Search Tool) search against the NCBI (National Center for Biotechnology Information) database. DNAsp (Rozas et al. 2017) was used to estimate haplotype diversity of all three species based on morphological assignment for each population. In order to visualize haplotype sharing between species, a parsimony haplotype network was created in PopART version 1.7 (Leigh and Bryant 2015) using the TCS method (Clement et al. 2000).

### Microsatellite DNA amplification and analysis

Eight microsatellite loci were amplified in *P. alabamensis, P. concinna*, and *P. floridana*. These microsatellites were originally developed by King and Julian (2004) who isolated 30 microsatellite loci in *P. floridana*. Eight of these microsatellites were later shown to amplify successfully in *P. alabamensis* (Hieb et al. 2011) and were used in our study. Each locus was run separately in 25ul reactions prepared using 5ul 5x GoTaq Flexi buffer, GoTaq Flexi DNA Polymerase 5u/ul (Promega), 0.5ul 25mM dNTPs, 2ul 25mM MgCl2, 1.2ul each of 10mM forward and reverse primers, 11.98ul H2O, and 3ul DNA extract. Thermal cycler conditions for amplification of all eight microsatellites were as follows: 94°C for 2 min, 35 cycles of 94°C for 45 s, 58°C for 45 s, 72°C for 1 min; and a final 5 min extension at 72°C. Fragment analysis of amplified products was performed by the DNA Analysis Facility at Yale University. Fragment lengths were scored manually using Peak Scanner Software Version 2.0 (Applied Biosystems, Carlsbad, Ca). For *P. floridana*, for the single population (Weeks Bay) with more than a few individuals, we only amplified a subset (27) of all the available individuals, while we amplified the microsatellite loci for all the other individuals of this species sampled elsewhere (Table 1).

**Table 1.** Sampling effort and number of individuals captured for each species (ID based on morphological assignment) across rivers. Sampling effort is displayed as the number of trap nights. A “trap night” is one trap set for one night. Numbers next to sampled watersheds correspond to numbers on the map in Figure 1. Numbers of individuals for each sex are indicated in parentheses as (Male, Female, Juvenile-unsexed). #Individuals per effort *100 refers only to *P. alabamensis* captures. * indicates donated samples. ** indicates roadkill collection.

| Sampled Watershed | Sampling Effort | *P. alabamensis* | #Individuals per effort *100 | *P. concinna* | *P. floridana* |
| --- | --- | --- | --- | --- | --- |
| (1) Biloxi River | 68 | 11 (8,1,2) | 16% | 39 (19,10,10) | 0 |
| (2) Pascagoula River | 50 | 18 (6,11,1) | 34% | 7 (4,3,0) | 0 |
| (3) Bayou La Batre | 10 | 1 (1,0,0) | 10% | 3 (3,0,0) | 1 (0,1,0) |
| (4) Fowl River | 48 | 5 (2,2,1) | 14% | 11 (6,5,0) | 2 (1,1,0) |
| (5) Dog River | 42 | 16 (7,9,0) | 38% | 9 (5,4,0) | 1 (0,1,0) |
| (6) Mobile-Tensaw Delta** | 52 | 24 (4,19,1) | 48% | 32 (13,17,2) | 1 (0,0,1) |
| (7) Weeks Bay | 106 | 18 (4,14,0) | 0.17% | 26 (16,10,0) | 68(23,45,0) |
| (8) Wolf River* | NA | 1 (0,1,0) | - | 0 | 0 |
| (9) Waifs* | NS | 2 (0,1,1) | - | 0 | 0 |
| Total | 376 | 96 |  | 127 | 73 |

Null alleles and allelic dropout were checked per each single population and across populations using MicroChecker (Van Oosterhout et al. 2004). Because null alleles can bias population structure analysis, FreeNA was used to calculate “uncorrected” and “corrected” (ENA correction, Chapuis & Estoup 2007) pairwise Fst (Chapuis & Estoup 2007) between river populations with N ≥ 5, between species and between STRUCTURE identified clusters (see below). Allelic diversity, presence of private alleles, observed and expected heterozygosities (H_O_, H_E_), and inbreeding coefficient (F_IS_) were assessed with the software Genetix v. 4.05 (Belkhir et al. 2004). Private alleles were considered for each population within each species (Petit et al. 2008) and for each species without distinction of populations. ARLEQUIN version 3.5.2.2 (Excoffier and Lischer, 2010) was used to calculate significance of Fst values, linkage disequilibrium between loci across all populations, and departure from Hardy-Weinberg equilibrium. BOTTLENECK v 1.2.02 (Cornuet & Luikart 1996) was used under all three mutational models available to detect signatures of historic bottlenecks within populations. The program Ne ESTIMATOR was used to infer breeding population size estimates for each river population (Do et al. 2014).

To identify patterns of genetic structure across the study area for *P. alabamensis*, we used the program STRUCTURE version 2.3.4 (Pritchard et al. 2000). We used the correlated allele frequency model with admixture to examine: all *Pseudemys* captured as a whole, *P. alabamensis* alone, and *P. concinna* alone. Since for *P. floridana* only a few individuals were found outside of the Weeks Bay system (Table 1), this species was not run independently of the others. STRUCTURE analysis consisted of ten independent runs for each K value (1-10) with a burn-in period of 100,000 followed by an additional 100,000 repetitions. In order to determine the best value of K (number of clusters) for each species, we used the ΔK statistic (Evanno et al. 2005) calculated using STRUCTURE HARVESTER (Earl 2012). STRUCTURE was also used to calculate the estimated membership coefficients Q for each individual in each cluster. Q indicates if each individual belongs to one or, if admixed, to several clusters.

## RESULTS

In total, 296 *Pseudemys* turtles were captured from water bodies known to be inhabited by *P. alabamensis* (Table 1). 96, 127, and 73 of these individuals were morphologically identified as *P. alabamensis, P. concinna*, and *P. floridana*, respectively. Despite many attempts, capture rates of *P. alabamensis* for some localities (Fowl River and Biloxi River) were low (Table 1), suggesting low population densities. One *P. alabamensis* individual was found in Wolf River, Mississippi, which is outside the currently recognized range of this species. Two potential hybrids between *P. alabamensis* and other *Pseudemys* species were identified in the field on the basis of morphological characteristics (Fig. 2). One of these individual’s, caught in Bayou La Batre, Alabama, appeared to be a *P. alabamensis* x *P. concinna* hybrid based on multiple morphological features including a strongly reduced jaw cusp, incomplete eye bars, and incomplete prefrontal arrow formed from the meeting of the sagittal head stripes with the supratemporal stripes. The other potential hybrid, captured in Dog River, resembled *P. peninsularis*, a non-native species but still possessed the identifying characteristics of *P. alabamensis. P. alabamensis* and *P. concinna* co-occurred in all the sampled rivers, while *P. floridana* mostly co-occurred with these other two species in Weeks Bay (Table 1).

In *P. alabamensis*, samples from the Biloxi River showed a skew toward males (Table 1). However, an abundance of hatchling *P. alabamensis* were observed in the area at the time of sampling (Moreno pers. obs.). For the Mobile-Tensaw Delta, our sampling included more females than males, as a large portion of our samples for this area came from road-kill individuals, which affects female turtles more than males (Marchand & Litvaitis 2004, Steen & Gibbs 2004). Overall, for *P. concinna*, more males than females were captured at all sites, except for the Mobile-Tensaw Delta.

### Mitochondrial DNA Analysis

A 587 bp fragment of the mtDNA control region was amplified from all 296 *Pseudemys* turtles sampled. Only 2 haplotypes were recovered for *P. alabamensis*: one haplotype (ARBT) was common among all sampled populations, while the other (Pen) was present only in a single individual from Dog River (Table 2, Fig. 3). BLAST search confirmed the common ARBT haplotype to be *P. alabamensis*, which was identical to a previously found haplotype (Jackson et al. 2012) (GenBank: GQ395751). The individual from Dog River with the Pen haplotype exhibited mixed morphological characteristics. This haplotype is four mutational steps from the ARBT *P. alabamensis* haplotype and matched *P. peninsularis* (GenBank: KC687235), a species that is normally only found on the Florida peninsula, indicating that this lone individual found in Dog River could be a hybrid. Out of 200 samples of *P. concinna* and *P. floridana*, 19 variable nucleotide positions -- including one insertion found in two individuals from Biloxi, Mississippi (haplotype = MissCon7) -- were identified, defining 22 haplotypes. One of these haplotypes was the ARBT of *P. alabamensis*, which was found in four individuals of *P. floridana* and two of *P. concinna*. Twelve haplotypes were unique to individuals morphologically identified as *P. concinna* (Con1, Con2, AlCon1, MissCon3, MissCon7, AlCon5, MissCon6, AlCon7, AlCon8, MissCon1, MissCon2, MissCon4), three haplotypes were unique to individuals morphologically identified as *P. floridana* (AlFlor3, AlFlor4, AlFlor5), and six haplotypes were shared between *P. concinna* and *P. floridana* (AlCon2, AlCon3, AlCon4, AlCon6, AlFlor1, AlFlor2) (Fig. 3). Of these individuals with shared haplotypes between species, 25 individuals morphologically identified *P. concinna* displayed *P. floridana* haplotypes and five individuals morphologically identified as *P. floridana* displayed *P. concinna* haplotypes. The star organization of the 12 haplotypes unique to *P. concinna* suggests a population expansion from the most represented haplotype (Con1) for this species. Haplotype diversity for *P. concinna* averaged 0.76 (range 0.53-0.86 among populations) and was 0.64 in the *P. floridana* Weeks Bay population (Table 2), which is the only population of this species with N>5. Haplotype sequences have been deposited to NCBI GenBank (see Data Accessibility section for accession numbers).

**Table 2.** Sample sizes and genetic diversity for each population of each species for the mitochondrial control region marker (mtDNA).

| Species | Sampled Watershed | N (mtDNA) | Number mtDNA Haplotypes | mtDNA Haplotype Diversity |
| --- | --- | --- | --- | --- |
| *P. alabamensis* | Weeks Bay | 18 | 1 | 0 |
|  | Mobile-Tensaw Delta | 24 | 1 | 0 |
|  | Dog River | 16 | 2 | 0.125 |
|  | Fowl River | 5 | 1 | 0 |
|  | Pascagoula River Delta | 17 | 1 | 0 |
|  | Biloxi River | 11 | 1 | 0 |
| *P. concinna* | Weeks Bay | 26 | 9 | 0.837 |
|  | Mobile-Tensaw Delta | 32 | 7 | 0.778 |
|  | Dog River | 9 | 5 | 0.861 |
|  | Fowl River | 11 | 5 | 0.818 |
|  | Pascagoula River Delta | 7 | 4 | 0.714 |
|  | Biloxi River | 37 | 6 | 0.53 |
| *P. floridana* | Weeks Bay | 68 | 9 | 0.637 |

**Figure 3.**
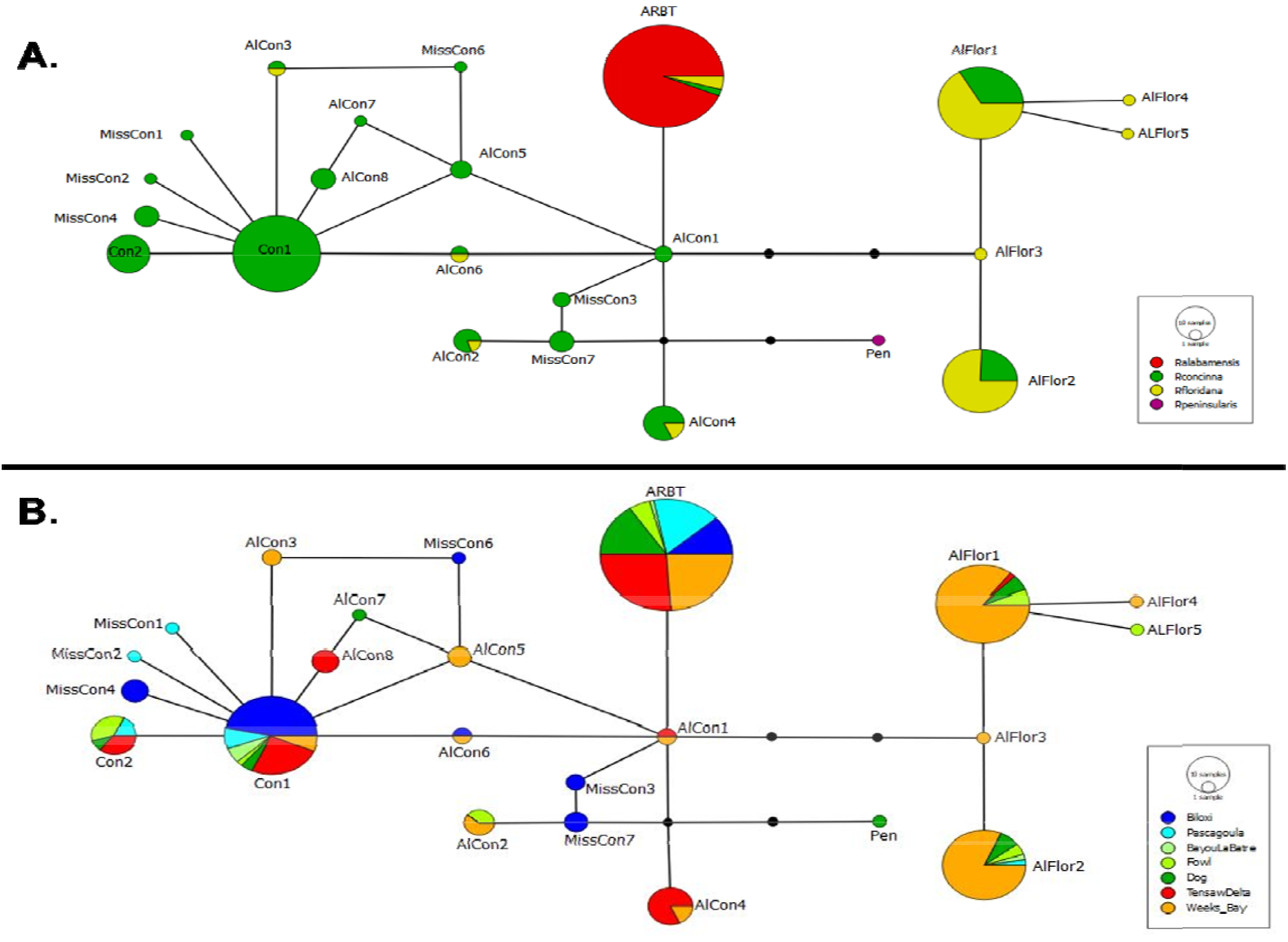
Haplotype networks produced from mitochondrial control region sequence data. **A**. Haplotype network showing the connectivity and haplotype sharing among species. **B**. Haplotype network showing haplotype distribution and sharing among sampling localities.

### Microsatellite

The eight microsatellite loci analyzed were polymorphic in all the species and populations, with the exception of one locus (D87) in one population (Fowl River) for *P. alabamensis*, and two loci (B91 in Pascagoula and D55 in two populations, Pascagoula and Fowl River) in *P. concinna*. The eight loci resulted in between 4-10 alleles each for *P. alabamensis*, 4-17 alleles each for *P. concinna*, and 3-12 alleles each for *P. floridana*. In *P. alabamensis*, private alleles were found exclusively in the Mobile Bay populations with the Mobile-Tensaw Delta possessing four private alleles, Weeks Bay three private alleles, and Dog River two private alleles (Table 3). In *P. concinna*, private alleles were found in all but the Fowl River population with Weeks Bay and Biloxi River possessing the most private alleles (nine and five, respectively) (Table 3). For each species, *P. alabamensis* had 14 private alleles in total, *P. concinna* 20, and *P. floridana* 5.

**Table 3.** Sample sizes and genetic diversity indices for each population (with N≥5) of each species for microsatellite data. N = sample size, NA = number of alleles, NU = number of private alleles, H_O_ and H_E_ = observed and expected heterozygosity, respectively, A = allelic diversity (Average number of alleles / locus), and F_IS_ = inbreeding coefficient

| Species | Location | N | NA | NU | He | Ho | A | $F_{IS}$ |
| --- | --- | --- | --- | --- | --- | --- | --- | --- |
| *P. alabamensis* | Biloxi River | 11 | 24 | - | 0.55 | 0.45 | 3.38 | 0.222 (0.015 - 0.301) |
|  | Pascagoula River | 17 | 26 | - | 0.47 | 0.39 | 3.63 | 0.194 (0.038 - 0.285) |
|  | Fowl River | 5 | 20 | - | 0.45 | 0.35 | 2.88 | 0.329 (-0.12 - 0.381) |
|  | Dog River | 16 | 32 | 2 | 0.55 | 0.48 | 4.63 | 0.157 (-0.011 - 0.254) |
|  | Mobile Tensaw Delta | 24 | 33 | 4 | 0.55 | 0.45 | 4.75 | 0.198 (0.067 - 0.279) |
|  | Weeks Bay | 18 | 30 | 3 | 0.54 | 0.47 | 4.37 | 0.147 (-0.031 - 0.257) |
| *P. concinna* | Biloxi River | 38 | 40 | 5 | 0.57 | 0.40 | 5 | 0.312 (0.218 - 0.376) |
|  | Pascagoula River | 7 | 24 | 1 | 0.45 | 0.45 | 3 | 0.074 (-0.20 - 0.133) |
|  | Fowl River | 11 | 31 | - | 0.47 | 0.42 | 3.88 | 0.112 (-0.119 - 0.204) |
|  | Dog River | 9 | 33 | 2 | 0.50 | 0.42 | 4.13 | 0.224 (-0.033 - 0.317) |
|  | Mobile Tensaw Delta | 20 | 42 | 4 | 0.56 | 0.46 | 5.25 | 0.206 (0.053 - 0.30) |
|  | Weeks Bay | 14 | 35 | 9 | 0.47 | 0.38 | 4.38 | 0.216 ( 0.057 - 0.277) |
| *P. floridana* | Weeks Bay | 27 | 46 | - | 0.51 | 0.41 | 5.75 | 0.122 (-0.002 - 0.21) |

Genetic diversity, as allelic diversity and heterozygosity, was generally low. Allelic diversity (A) in populations with N ≥5 ranged from low (2.88) to moderate (4.75) in *P. alabamensis* (mean 3.94), from 3-5 (mean 4.27) in *P. concinna*, and was relatively higher (5.75) in the single *P. floridana* population found in Weeks Bay (Table 3). Observed heterozygosity (H_O_) ranged from 0.35 - 0.48 in *P. alabamensis*, 0.38 - 0.45 in *P. concinna*, and was 0.41 in *P. floridana*. Expected heterozygosity (H_E_) ranged from 0.45 - 0.55 in *P. alabamensis*, 0.45 - 0.57 in *P. concinna*, and was 0.51 in *P. floridana*. All populations for all species, except for Pascagoula in *P. concinna*, show an excess of homozygosity with H_O_ having much lower values than H_E_. In *P. alabamensis*, bottleneck analysis identified one significant occurrence (P < 0.05) for the Pascagoula population (N=17) under the Stepwise Mutation Model. Ne estimates show support for slow breeding population sizes (Ne < 30) in the Biloxi River and Fowl River populations of *P. alabamensis* (Table 4). Consequently, inbreeding was observed for these two populations with F_IS_ values of 0.22 and 0.33, respectively (Table 3). Ne ESTIMATOR found little evidence of low breeding population sizes in *P. concinna* or *P. floridana* (Table 4).

**Table 4.**
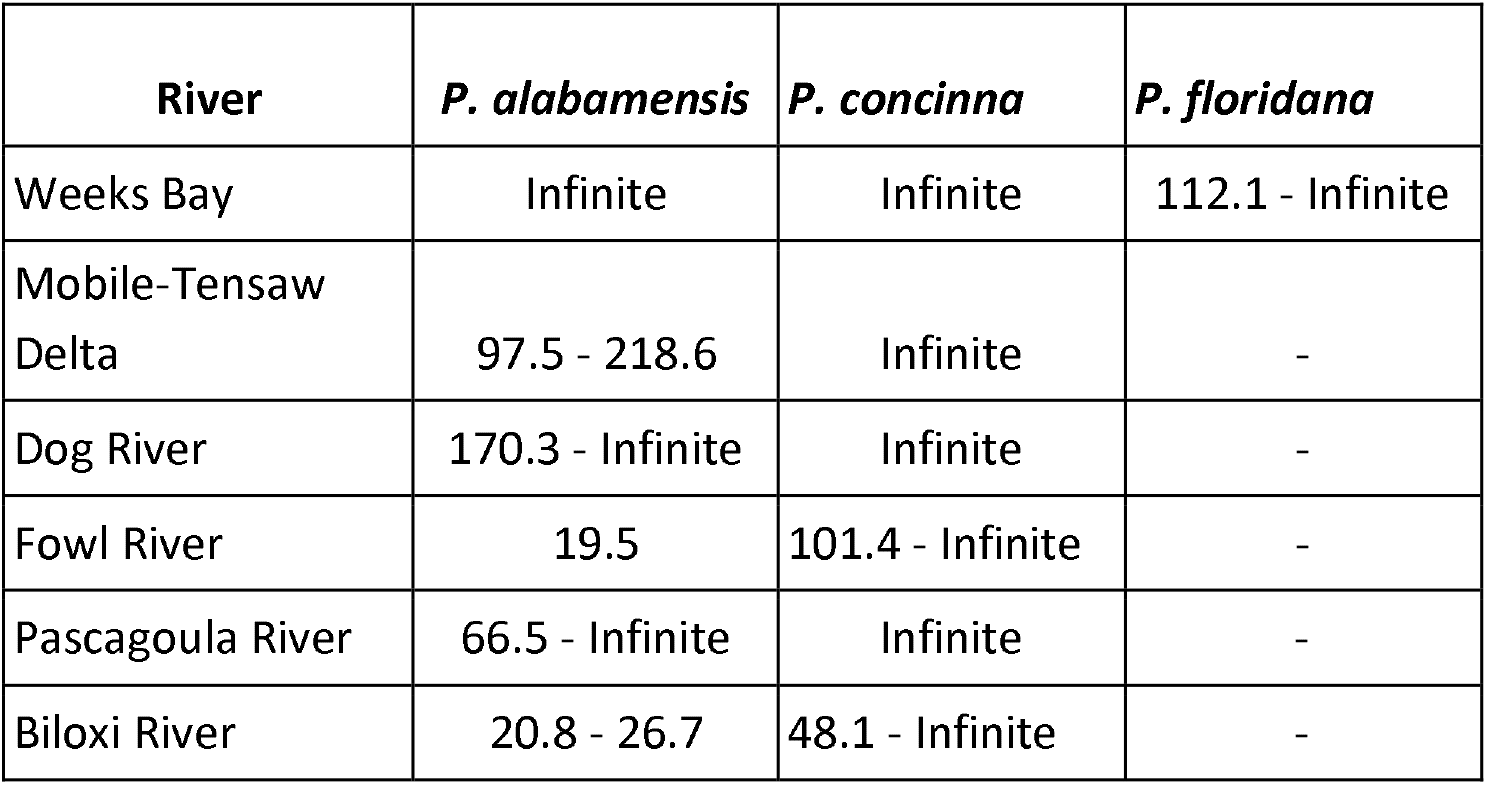
Ne ESTIMATOR breeding population size estimates.

| **River** | ***P. alabamensis*** | ***P. concinna*** | ***P. floridana*** |
| --- | --- | --- | --- |
| Weeks Bay | Infinite | Infinite | 112.1 - Infinite |
| Mobile-Tensaw<br>Delta | 97.5 - 218.6 | Infinite | - |
| Dog River | 170.3 - Infinite | Infinite | - |
| Fowl River | 19.5 | 101.4 - Infinite | - |
| Pascagoula River | 66.5 - Infinite | Infinite | - |
| Biloxi River | 20.8 - 26.7 | 48.1 - Infinite | - |

Two alleles, B21 and D79, showed evidence of null alleles in all three species for half or more of the sampled populations, with the Biloxi population especially affected by the presence of null alleles in *P. concinna*. Overall, null alleles were recovered in 23 out of the 96 combinations of loci x populations x species (8 loci, 6 populations, 2 species with various populations). F-tests run on corrected and not corrected Fst values obtained using FreeNA indicate that the presence of null alleles does not affect Fst estimates (p-value >0.05 for each species comparison). Therefore, all microsatellite loci were used in subsequent analyses. No loci showed significant linkage disequilibrium (p < 0.01) across populations providing evidence of independent segregation of loci used. All populations of *P. alabamensis* and all but one of *P. concinna* show departure from Hardy-Weinberg Equilibrium at the loci D79, most likely as a result of the null allele and higher homozygosity levels. Among all populations, *P. concinna* from Biloxi possesses the most significant departures at five of the eight loci, with lower than expected heterozygosity. Similarly, the single *P. floridana* population with N>5 displayed significant departure from Hardy-Weinberg equilibrium in three loci. Within species, Fst values among populations ranged between 0-0.28 and 0-0.19 for *P. alabamensis* and *P. concinna*, respectively. Populations from Pascagoula (Mississippi) for both *P. alabamensis* and *P. concinna* were the most distinct (Fst values >0.17) from counterpart populations in Alabama (Table 5). Pairwise Fst values calculated for *P. alabamensis* vs. *P. concinna* and *P. floridana* were 0.106 and 0.132, respectively, while Fst between *P. concinna* and *P. floridana* was found to be low (0.065), likely as a result of admixture between the two.

**Table 5.** Fst pairwise values based on microsatellite data for populations of *P. alabamensis* (bottom left), and *P. concinna* on the top right axis. Bold values are significant at P<0.05.

|  | Weeks Bay | Mobile-Tensaw Delta | Dog River | Fowl River | Pascagoula River | Biloxi River |
| --- | --- | --- | --- | --- | --- | --- |
| Weeks Bay | - | **0.103** | **0.110** | **0.123** | **0.186** | **0.102** |
| Mobile-Tensaw | **0.025** | - | **0.035** | 0.021 | **0.147** | **0.046** |
| Dog River | **0.027** | 0.000 | - | 0.001 | **0.148** | **0.060** |
| Fowl River | **0.145** | **0.093** | **0.068** | - | **0.193** | **0.045** |
| Pascagoula River | **0.222** | **0.225** | **0.172** | **0.282** | - | **0.138** |
| Biloxi River | **0.100** | **0.077** | **0.046** | **0.125** | **0.095** | - |

STRUCTURE analysis of all *Pseudemys* species considered in this work identified an optimum clustering of K=2 with evidence of some admixture (Fig. 4). The two clusters corresponded to *P. alabamensis* and *P. concinna*/*P. floridana*, respectively. When the clustering analysis was performed only on *P. alabamensis*, optimum clustering was also K=2 corresponding to Mississippi and Alabama populations. The analysis repeated only on *P. concinna* found an optimum clustering level of K=3 corresponding to (1) the Biloxi River population, (2) Fowl River, Dog River, and Tensaw Delta populations, and (3) the Pascagoula River and Weeks Bay populations. Fst of *P. concinna* clusters were generally low and follow the cluster numbers listed above: cluster 1 vs 2 = 0.043, cluster 1 vs 3 = 0.078, cluster 2 vs 3 = 0.072. (see Data Accessibility section for files with microsatellite allele scoring - available after manuscript acceptance as Supplementary Material S1).

**Figure 4.**
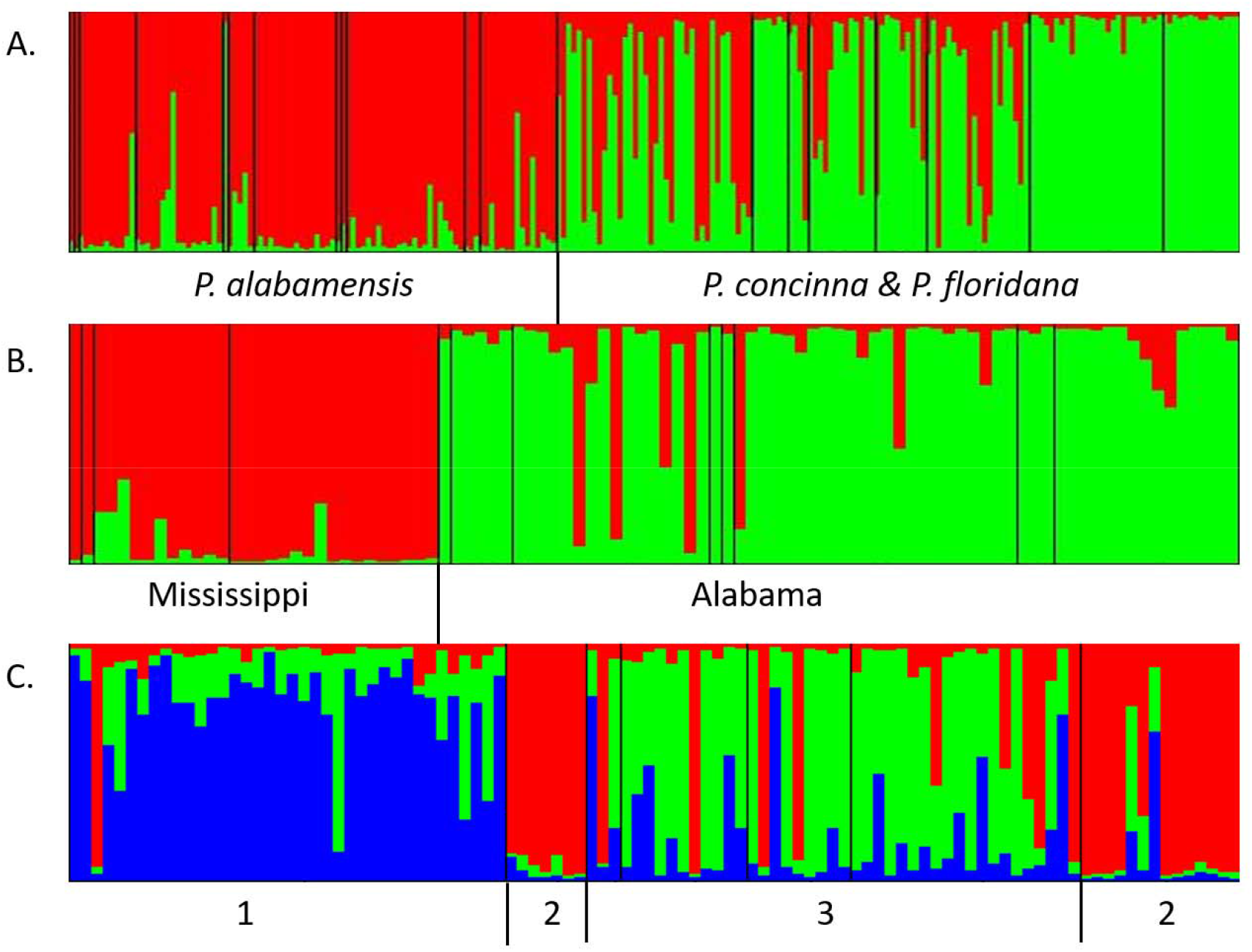
Structure graphs created from microsatellite data consisting of 8 loci. **A**. Structure graph of all turtles sequenced showing clusters of *P. alabamensis* and next to sympatric cooter species. **B**. Structure graph of *P. alabamensis* showing Mississippi and Alabama clusters. **C**. Structure graph of *P. concinna*. Subdivisions of *P. concinna* structure graph as follows: 1. Biloxi population, 2. Pascagoula and Weeks Bay population, 3. Populations from Mobile County, Alabama.

### Hybridization

Of the 96 morphologically identified *P. alabamensis* samples, two individuals were considered to be potential hybrids based on mixed morphological characteristics and presence of reduced *P. alabamensis* identifying characteristics. One of these individuals - an individual from Dog River - possessed a *P. peninsularis* mtDNA haplotype, seven microsatellite loci with alleles matching *P. alabamensis* alleles, and one microsatellite locus possessing an allele not found in any other individual of the species studied here. The other individual (from Bayou La Batre), despite having the *P. alabamensis* mtDNA haplotype (ARBT), was recovered with the cooter species (*P. concinna* and *P. floridana*) in the STRUCTURE analysis based on microsatellite loci. Of the 102 *P. concinna* and 30 *P. floridana* with both mtDNA data and microsatellite data, 27 (26.5%) of *P. concinna* and 6 (20%) of *P. floridana* possessed conflicting species assignments between the two marker types. Two morphologically identified *P. concinna* and four *P. floridana* individuals possessed the *P. alabamensis* haplotype (ARBT), although for *P. floridana* they did not group with *P. alabamensis* in the STRUCTURE analysis based on microsatellites. One of these four individuals possessed a bright red plastron, a characteristic not present in *P. floridana*, which typically possess plain yellow plastrons; however no other potential *P. alabamensis* morphological characteristics were seen in these six individuals. Five of the six cooters that displayed the *P. alabamensis* haplotype were found in the rivers of Weeks Bay, while the sixth was found in the Mobile Tensaw Delta.

Based on microsatellite data, Fst between *P. alabamensis* and each of the other two sympatric *Pseudemys* species is lower (from *P. concinna* and *P. floridana* were 0.106 and 0.132, respectively) than Fst observed within *P. alabamensis* from Pascagoula versus the populations in Alabama (Fst ranging from 0.17 to 0.28, Table 5), further supporting the occurrence of hybridization between species. Between *P. alabamensis* and *P. concinna*, three of the alleles private to populations within a single species were found in the other species. Sharing of private alleles may be an indication of admixture and introgression. One private allele from locus B91 that was only found in the Dog River population of *P. alabamensis* was also found in the Biloxi River population of *P. concinna* (frequency of the allele in Biloxi = 0.026). One private allele from locus D121 that was only found in the Weeks Bay population of *P. alabamensis* was also found to be a common allele in *P. concinna* (frequency of the allele in *P. concinna* reached 0.278 in the Mobile-Tensaw Delta population). And one allele from locus D28 that was a private allele in the Biloxi *P. concinna* population was also found in the neighboring Pascagoula population of *P. alabamensis* (frequency of the allele in *P. alabamensis* in Pascagoula = 0.027).

Hybridization appears to occur at a higher rate between *P. concinna* and *P. floridana*. Fst between the two species is 0.065, much lower than between *P. alabamensis* and any of these two species (see above) and even within *P. alabamensis*. The *P. floridana* population with more than 5 individuals possessed 6 of the alleles that were found to be private alleles within *P. concinna* populations and 1 allele that was considered a private allele within a *P. alabamensis* population.

When examining species assignment and admixture by STRUCTURE, we found three individuals of *P. alabamensis* to be assigned to *P. concinna* (one from Bayou La Batre with Q = 0.95 was assigned to *P. concinna*/*P. floridana*, one from Fowl River with Q = 0.68, and one from Weeks Bay with Q = 0.66 to *P. concinna* and 0.77 to *P. floridana*). Another two individuals morphologically ID as *P. alabamensis*, one from Pascagoula and one from Fowl River, showed admixture with mixed assignment between *P. alabamensis* and *P. concinna*. Signs of hybridization with *P. alabamensis* were also found in individuals morphologically identified as *P. concinna*. Out of the 102 individuals morphologically ID as *P. concinna*, 12 individuals (∼12%) were assigned to *P. alabamensis* with Q > 0.7, and another 14% showed mixed assignment between the two species. Across all the populations, the Biloxi river was the locality where many individuals morphologically identified as *P. concinna* were assigned to *P. alabamensis* on the basis of microsatellite data. We also found three individuals (out of 30, 10%) morphologically identified as *P. floridana* showing evidence of admixture (all the Q values can be found in the Supplementary Material S2). Finally, out of all individuals of *P. concinna* and *P. floridana*, 42 of 103 (40.77%) *P. concinna* were either assigned to *P. floridana* or showed admixture and 14 of 30 (46.66%) *P. floridana* were also either assigned to *P. concinna* or showed admixture.

## Discussion

In this study, we assessed the genetic diversity, population structure, and potential hybridization of the endangered *P. alabamensis* and co-occurring congeneric species. While previous studies have also addressed some of these questions (Jackson et al. 2012, Heib et al. 2014), the sample sizes, distribution range of sampled populations, and/or genetic markers were limited. In our study, we used both mitochondrial and microsatellite markers to analyze *P. alabamensis* from seven rivers throughout the entire narrow range of the species. Using mitochondrial DNA, we found no genetic differentiation within or among populations of *P. alabamensis* due to a complete lack of mtDNA variation. Low levels of mitochondrial diversity is not uncommon in turtles that are of conservation concern (e.g., Rosenbaum et al. 2007, Vargas-Ramírez et al. 2007). However, differently from what has been observed in other endangered species, only one haplotype was found across 96 individuals from the entire distribution range of *P. alabamensis*. A comparable lack of mitochondrial diversity to *P. alabamensis* has also been noted in a related species, *Pseudemys* gorzugi (Bailey et al. 2008). *P. gorzugi* also inhabits a restricted range, although larger, being found only in the Rio Grande and Pecos Rivers in North America. While mtDNA is used extensively for phylogeographic studies because of its relatively high mutation rate, maternal inheritance, and ease of amplification, it is known to have a slow rate of evolution in turtles (Amato et al. 1997; Avise 2000; King and Julian 2004; see also Lourenço et al. 2013). *P. alabamensis* displayed no genetic variation across populations at the mtDNA control region; conversely, *P. concinna* and *P. floridana* showed a higher degree of genetic variation within the same river populations. This may be indicative of the larger population sizes and may reflect the greater overall distribution range compared to *P. alabamensis. P. concinna* populations in the area are likely connected to larger populations occurring in northern Alabama and Mississippi through the larger rivers of the Mobile-Tensaw Delta and Pascagoula Delta watersheds. Individuals dispersing from the northern populations may contribute to the genetic variation of the smaller isolated coastal populations.

Microsatellite data also indicate low genetic diversity for *P. alabamensis* with overall lower allelic diversity than the other two sympatric congenerics and a lower than expected heterozygosity. Signs of inbreeding were observed in two populations: Fowl River and Biloxi Rivers. Biloxi showed signs of inbreeding also for *P. concinna*, most likely the result of low population sizes for both species at this site (the estimated breeding population for *P. alabamensis* at Biloxi was in fact low; see also hybridization part below). Despite the overall low genetic diversity observed in *P. alabamensis*, microsatellite data support genetic differentiation between Mississippi and Alabama populations of this species, in agreement with slight morphological differences previously observed between these areas (Leary et al. 2003). This observed genetic differentiation may be due to the non-connectivity of the populations due to the large distance between the mouth of the Pascagoula River Delta and the Alabama populations. For freshwater species distributed across the Gulf of Mexico, several riverine systems have been found to act as barriers to gene flow (Soltis et al. 2006) including the Pascagoula River (e.g., Dugo et al. 2004, Ennen et al. 2010).

We found no structure among populations of *P. alabamensis* that make up the Mobile Bay (populations 3-7 in Fig. 1). This may be due to potential migration of individuals between these populations due to the lower salinity of the Mobile Bay compared to the Mississippi Sound. Movement of individuals across populations, including towards the lower part of the Mobile-Tensaw Delta may be permitted by the fact that *Pseudemys* species have been reported to possess some level of tolerance to brackish water (Agha et al. 2018). This is further supported by the presence of barnacles on the shells of some individuals in our study indicating exposure to higher salinity waters (Moreno pers. obs.). The presence of multiple alleles that are found in all major Alabama populations, but not in Mississippi populations also suggests the occurrence of gene flow among the Alabama populations.

We observed admixture between the three species. Individuals of *P. concinna* and *P. floridana* in the region can be difficult to tell apart due to hybridization between the two (Mount 1975, Spinks et al. 2013). We found that even for individuals which could be confidently assigned to one or the other species based on morphological characteristics, haplotype sharing and mixed or “different from morphological” assignment based on microsatellite data was observed between species. Specifically, based on microsatellite data, more than 40% of the individuals that were morphologically identified as *P. concinna* had mixed assignment with *P. floridana* and vice versa. Haplotype sharing is also seen to a lesser degree between *P. alabamensis* and *P. concinna*/*P. floridana* individuals and it is confirmed by microsatellite data. To our knowledge, these data represent the first published evidence of hybridization between *P. alabamensis* and sympatric *Pseudemys* species. In all of these cases of haplotype sharing, animals were morphologically identified as *P. concinna* or *P. floridana* but had the *P. alabamensis* mtDNA haplotype, suggesting that hybridization in *P. alabamensis* may be largely driven by males of *P. concinna* and *P. floridana* breeding with female *P. alabamensis*. In Weeks Bay and in the Mobile-Tensaw Delta, where we found instances of haplotype sharing among species, we sampled an excess of female versus male *P. alabamensis* (Table 1). Hybridization of *P. alabamensis* with congeneric species across its distribution range may overall be driven by decreased opportunities to find mates of the same species, from potentially a skewed sex ratio in Weeks Bay and Mobile-Tensaw Delta to low breeding population sizes in Biloxi, Bayou La Batre, and Fowl River (Tables 1, 3, and 4). In Biloxi for example, we found many individuals morphologically identified as *P. concinna*, but genetically assigned to *P. alabamensis*. in Bayou La Batre, we also found a single specimen morphologically identified as *P. alabamensis* possessing strongly reduced redbelly (a characteristic of *P. alabamensis*). This individual grouped with cooter species in a STRUCTURE analysis, but had the mtDNA haplotype of *P. alabamensis* suggesting a possible *P. concinna* x *P. alabamensis* hybrid origin. Finally, based on mtDNA data, in Dog River we found only one female that was morphologically identified as *P. alabamensis* that possessed a *P. peninsularis* haplotype. The home range of *P. peninsularis* is isolated to the Florida peninsula and is not native to the range of *P. alabamensis*. It is possible that this individual represents a *P. peninsularis* x *P. alabamensis* hybrid offspring of a female *P. peninsularis* that was released into Dog River and bred with native *P. alabamensis*. Overall, based on our results, *P. alabamensis* is experiencing some level of admixture with congeneric co-occurring species across its entire and restricted distribution range. Hybridization for species of conservation concern with limited population sizes is a well known phenomena (see for example Chattopadhyay et al. 2018 and references therein), and it presents a challenge for management and conservation actions (Mallet 2005, Allendorf et al. 2005, Wayne & Shaffer 2016).

Our study represents the first genetic study supporting the endangered status of *P. alabamensis* throughout its range and providing evidence that the Mississippi and Alabama populations of *P. alabamensis* are genetically different. Despite this genetic distinction, the overall low amount of genetic diversity observed at the mitochondrial and nuclear (microsatellite) levels in *P. alabamensis*, the severely limited geographic range of this species, and the occurrence of hybridization throughout its distribution, require the urgent development of targeted conservation actions. Local monitoring activities to ensure habitat protection of the few sites where the species occurs, maintenance of nesting sites, assessment of recruitment throughout the species’ range, and monitoring of population sizes should be developed for this species. Our results also identify populations of higher conservation concern, because of low population sizes and consequent inbreeding and hybridization: Bayou La Batre, Biloxi, Weeks Bay, and Fowl River. Furthermore, considering the observed genetic distinction of populations from Alabama and Mississippi, specific management actions should be developed to preserve their uniqueness. This also includes search for additional unknown branches of these main riverine systems where the species could occur. Finally, as climate change will strongly affect coastal areas and wetlands on the Gulf of Mexico (Mulholland et al. 1997, Scavia et al. 2002, Anderson et al. 2014), and influence the geographic range of species (e.g., Garroway et al. 2010), the imperiled status of *P. alabamensis* may further worsen due to changes in salinity of the water in its habitat, effects on the vegetation on which this species feed, and potentially increased hybridization. It is therefore imperative that measures to prevent progressive declining of populations and mitigate current and future effects of climate change on *P. alabamensis* are considered and developed rapidly.

## Supplementary Material

### Supplementary Material S1

Microsatellite genotyping for each individual and each locus (to be provided after manuscript acceptance).

### Supplementary Material S2

Q assignment value for each individual with respect to each different species and *P. alabamensis* clusters (Alabama vs. Mississippi). To be provided after manuscript acceptance.

## Data Accessibility Statement

Haplotype sequence data has been deposited in the NCBI GenBank (accession numbers: MZ966274-MZ966297). Microsatellite genotyping for each individual and each locus will be provided as Supplementary Material S1 after manuscript acceptance.

## Competing Interests Statement

None of the authors have competing interests.

## Author contributions

Y.C., S.G, and D.N. conceived the study. N.M., A.H., K.B., and E.M. collected samples. N.M. performed lab work. N.M. performed the analyses, N.M., S.G., and Y.C. wrote the manuscript.

## Acknowledgments

We are thankful to Skyler Kerr, Dominika Houserova, Robin Loyd, and Nathan Katlein for help with field collections of turtles. We acknowledge the Central Mississippi Turtle Rescue and Dauphin Island Aquarium for providing waif samples of *P. alabamensis*. We are also grateful to Danielle Edwards and Guillermo Velo-Anton for feedback on some of the analyses performed in this study.

## Notes

### Competing Interest Statement

The authors have declared no competing interest.

